# Feedback adaptation to unpredictable force fields in 250ms

**DOI:** 10.1101/773002

**Authors:** Frédéric Crevecoeur, James Mathew, Marie Bastin, Philippe Lefevre

## Abstract

Motor learning and adaptation are important functions of the nervous system. Classical studies have characterized how humans adapt to changes in the environment during tasks such as reaching, and have documented improvements in behavior across movements. Yet little is known about how quickly the nervous system adapts to such disturbances. In particular, recent work has suggested that adaptation could be sufficiently fast to alter the control strategies of an ongoing movement. To further address the possibility that learning occurred within a single movement, we designed a series of human reaching experiments to extract in muscles recordings the latency of feedback adaptation. Our results confirmed that participants adapted their feedback responses to unanticipated force fields applied randomly. In addition, our analyses revealed that the feedback response was specifically and finely tuned to the ongoing perturbation not only across trials with the same force field, but also across different kinds of force fields. Finally, changes in muscle activity consistent with feedback adaptation occurred in about 250ms following reach onset. We submit this estimate as the latency of motor adaptation in the nervous system.

## Introduction

Humans and other animals can adapt motor patterns to counter predictable disturbances across a broad range of contexts, including reaching, locomotion, and eye movements (Shadmehr et al., 2010; Wolpert et al., 2011; Roemmich and Bastian, 2018). A central question in movement neuroscience is to identify the time scales at which this process can influence behavior. In the context of reaching movements, standard learning paradigms have focused on trial-by-trial learning, such that changes in behavior were documented by contrasting early and late motor performances, often separated by minutes to hours, or equivalently by hundreds of trials (Lackner and DiZio, 1994; Shadmehr and Mussa-Ivaldi, 1994; Singh and Scott, 2003; Smith et al., 2006; Wagner and Smith, 2008). Thus, a clear benefit of motor adaptation is to improve behavior over these timescales, which is of prime importance for instance when we deal with a new tool or environment. Clearly the associated neural mechanism must also be beneficial for adaptation to changes occurring over slower time scales such as development and long-term skill acquisition (Dayan and Cohen, 2011).

Besides the improvement of behavior over medium to long timescales, previous studies also indicated that motor learning could be very fast. The presence of rapid adaptation was previously established by observing after effects induced by a single movement (Sing et al., 2013). Likewise, unlearning was documented after a single catch trial when a force field was unexpectedly turned off (Thoroughman and Shadmehr, 2000). Our previous study showed that the timescale of motor learning could be even faster. Indeed, it was documented that healthy volunteers could produce adapted feedback responses to the unanticipated force field perturbations during reaching, and after effects were evoked within an ongoing sequence of movements in less than 500ms when participants were instructed to stop at a via-point (Crevecoeur et al., 2018).

These latter results contrasted with standard models of sensorimotor learning (Thoroughman and Shadmehr, 2000; Baddeley et al., 2003; Smith et al., 2006; Kording et al., 2007), which included multiple timescales but nevertheless assumed that each movement was controlled with a fixed representation. The existence of after effects in less than 500ms challenged this view, as it reveals that adaptation was potentially fast enough to influence an ongoing movement. Thus, motor adaptation could not only support learning across trials, but also complement online feedback control.

Evidence for online adaptation was interpreted in the context of adaptive control (Bitmead et al., 1990): a least-square learning algorithm coupled with a stat-feedback controller. This technique is based on standard state-feedback control models that successfully capture humans’ continuous and task-dependent adjustments of voluntary movements (Todorov and Jordan, 2002; Diedrichsen, 2007; Liu and Todorov, 2007), as well as feedback responses to mechanical perturbations (Scott, 2016; Crevecoeur and Kurtzer, 2018). Intuitively, the state feedback controller in the nervous system can be viewed as a parameterized control loop, and the goal of adaptive control is to tune this loop in real time by continuously tracking the model parameters (and errors). This model captured both adjustments of control during un-anticipated perturbations, and the standard single rate trial-by-trial learning observed across a few trials (Crevecoeur et al., 2018).

To gain further insight into the timescales of motor adaptation in the brain, we designed this study to address the following key questions: first we sought to reproduce previous findings of adaptation to unpredictable disturbances, and measure precisely the latency of adaptive changes in control from muscle recordings. Second, we sought to test a surprising prediction of the theoretical framework of adaptive control: if the nervous system tracks model parameters in real time, then, in principle, it should be possible to handle simultaneously force fields not only of different directions (clockwise or counterclockwise), but also of different kinds (i.e. with different force components). First, our results showed that feedback responses to unanticipated perturbations became tuned to the force field within ~250ms of movement onset. Second, we found that humans were indeed able to produce adapted and specific feedback responses to different force fields randomly applied as catch trials. Our results confirmed the existence of very fast adaptation of feedback control during movements and provide an estimate of ~250ms for the latency of motor adaptation.

## Methods

### Experiments

A total of 44 healthy volunteers were involved in this study. Participants provided written informed consent and the procedures were approved by the Ethics Committee at the host institution (UCLouvain, Belgium). Eighteen participants performed the first experiment, another group of 18 participants performed the second experiment, and the rest (n=8) performed the control experiment. The data of the control experiment was published in our previous study and was re-used here to underline the similarities between feedback adaptation and standard trial-by-trial learning (Crevecoeur et al., 2018)

In all experiments, participants grasped the handle of a robotic arm (KINARM, BKIN Technologies, Kingston, ON, Canada), and were instructed to perform visually guided reaching movements towards a virtual target. Each trial ran through as follows. Participants had to wait in the home target (a filled circle with radius 0.6cm) for a random period uniformly distributed between 2s and 4s. The goal was also displayed as a circle located 15cm ahead of the start. After the random period, the cue was delivered to initiate the movement by filling the goal target (Fig. 1a). Participants had between 600ms and 800ms (including reaction time) to reach the goal and stabilize in it during at least 1s. Information about the time window was provided as follows: when participants reached the goal too soon, it turned back to an open circle. When they reached it too late, it remained red. When they reached it within the desired time window, it became green and a score displayed on the screen was incremented. The scores and feedback about timing were provided to encourage consistent movement times, but all trials were included in the dataset. The grand average success rate was 70±12% for Experiment 1, and 76±10% for Experiment 2. In all cases, direct vision of the arm and hand was blocked but the cursor aligned to the handle was always visible. These procedures were identical across the three experiments, which only varied by the frequency and nature of mechanical perturbations applied during movements.

**Figure 1.**
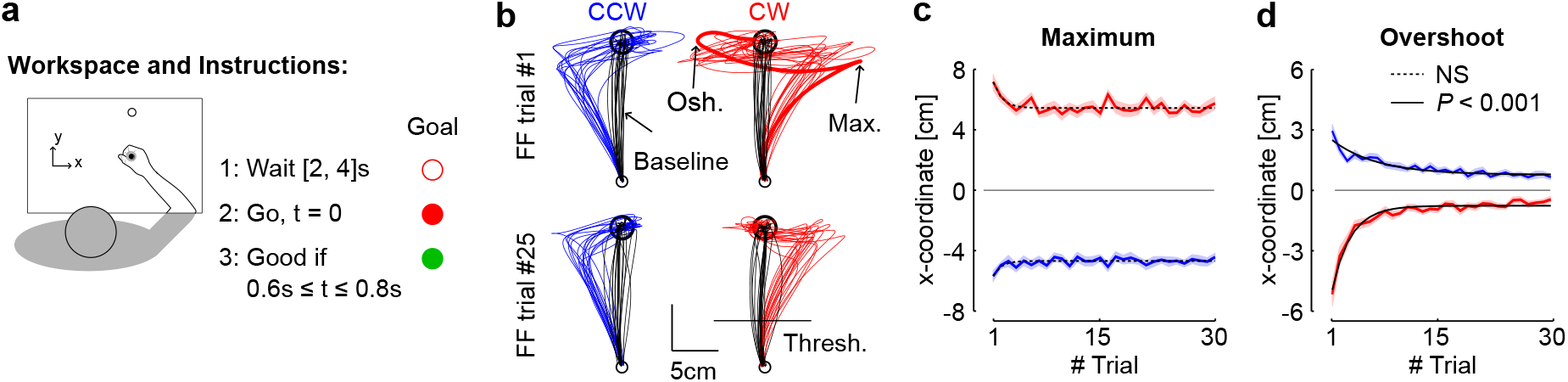
**a.** Illustration of the workspace and task. Participants were instructed to perform forward reaching movements towards a visual target. An open goal target was presented for a random delay uniformly distributed between 2s and 4s before it was filled in. The cue to reach the target was given by filling in the goal in red. The goal was turned red if the time between the go signal and the stabilization in the target was comprised between 0.6s and 0.8s. **b.** Hand paths from the first force field trials (top) and from the trial #25 selected for illustration (bottom) from each participant (n=18). Counter-clockwise and clockwise perturbations are depicted in blue and red, respectively. The black traces illustrate for each panel baseline trials selected randomly (1 baseline trial per participant). **c.** Maximum displacement in the direction of the force field. The dashed trace illustrate that the exponential fit did not reveal any significant curvature across force field trials (P>0.05). **d.** Maximum target overshoot in the direction opposite to the force field. Solid traces revealed strongly significant exponential decay across trials (P<0.001).

#### Experiment 1

This experiment was designed to reproduce previous results on adaptation of feedback responses to unpredictable perturbations (Crevecoeur et al., 2018), and to measure the moment within a trial when the muscle activity started to show feedback tuning corresponding to the force field. Participants performed six blocks of 60 trials, composed of unperturbed trials (baseline) and force field trials. The *x* and *y* coordinates corresponded to lateral and forward directions, respectively (Fig. 1a). In this experiment, the force field was defined as a lateral force proportional to forward velocity: 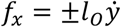, with *l*_*o*_ = ±13 Nsm^−1^ (the subscript “*O*” refers to the Orthogonal force field). There were 5 force field trials per block and per direction (counter-clockwise and clockwise), which corresponded to a frequency of perturbation trials of 1/6, and a total of 30 force field trials for each perturbation direction. The sequence of trials was randomized within each block, such that the occurrence and direction of the perturbations were unpredictable.

#### Experiment 2

The purpose of this experiment was to test further the hypothesis of online adaptive control by alternating different kinds of force fields, which in theory could be handled by online tracking of model errors (Bitmead et al., 1990). To investigate this, we performed an experiment similar to Experiment 1, with the addition of curl force field trials randomly interspersed between unperturbed and orthogonal force field trials. The orthogonal force field was identical to Experiment 1. For the curl field, both forward and lateral velocities were mapped onto lateral and forward perturbation forces with opposite signs, respectively: 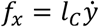, and 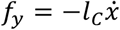 with *l*_*c*_ = ±15 Nsm^−1^ (the subscript “*C*” refers to the Curl field). There were 5 perturbation trials per force field (orthogonal and curl) and direction (clockwise and counterclockwise), summing to a total of 20 perturbations per block presented in a random sequence. As in Experiment 1, participants performed six blocks of 60 trials, composed of 40 baseline trials and 20 perturbation trials (perturbation frequency: 1/3).

#### Control experiment

In this experiment we were interested to measure participants’ behavior in a fully predictable context corresponding to a standard adaptation task. Participants performed a series of baseline trials for training, followed by 180 force field trials (orthogonal force field, CW or CCW for the entire series), followed by another series of 180 force field trials in the opposite direction for the entire series. The two series were separated by 20 baseline trials to induce washout between the two adaptation phases. We re-used previously published data for this experiment and refer to (Crevecoeur et al., 2018) for complementary descriptions of the results.

### Data Collection and Analysis

The 2-dimensional coordinate of the cursor aligned to the robotic handle, and the forces at the interface between the participants’ hand and the handle were sampled at 1kHz, and digitally low-pass filtered with a fourth-order dual-pass Butterworth filter with cut-off frequency of 50Hz. Velocity signals were obtained from numerical differentiation of position signals (4^th^ order, finite difference algorithm). We collected the activities of two of the main muscles recruited when performing lateral corrections against the perturbations used in our experiment: Pectoralis Major (shoulder flexor) and Posterior Deltoid (shoulder extensor). Muscles samples were recorded with surface electrodes for Experiments 1 and 2 (Bagnoli Desktop System, Delsys, Boston, MA, US). EMG signals were collected at 1kHz, digitally band-pass filtered (4^th^ order dual pass: [10, 400] Hz), and rectified.

Two events were used as timing references. First, reach onset was defined as the moment when the cursor aligned to the handle exited the home target. Second, we used a position threshold located at 1/3 of the distance between the home and goal targets to re-align the EMG traces offline. The crossing of this position threshold approximately coincided with the peak forward velocity, which allowed reducing the trial-to-trial variability in EMG recordings. Similar conclusions were obtained when all analyses were performed based on traces realigned with respect to reach onset.

Exponential fits were used to quantify the presence of learning on several parameters, including the maximum lateral hand displacement, and maximum target overshoot for Experiment 1. The quantification of learning from Experiment 2 was based on exponential fits of the path length computed as the time integral of hand speed. We fitted the exponential functions to the raw data from each participant as a function of the trial index, and assessed whether the 99.9% confidence interval for the parameter responsible for the curvature of the fit included or not the value of 0 (*P*<0.001). Variability across participants was illustrated on hand trajectories by calculating the dispersion ellipses based on singular value decomposition of the covariance matrices at different time steps evenly spaced.

We measured both the onset of changes in EMG responsible for changes in behavior across early and late force field trials, as well as the onset of changes in EMG across force fields from Experiment 2. To contrast early and late trials, EMG data was averaged for each participant across the first four and last four trials. To contrast the feedback responses to orthogonal and curl fields in Experiment 2, EMG data was averaged across the last 15 trials of each kind of force field. EMG averages were then collapsed into a 30ms wide (centered) sliding window, and sliding comparisons across time were performed with paired t-tests. We searched in the time series of *P*-values the moment of strongest statistical difference across populations of EMG data (*P*<0.005), and then went back in time until the threshold of *P*<0.05 was crossed. On the one hand, this test could identify early differences since it included data from −15ms to +15ms relative to the center of the bin, but on the other hand, we kept the threshold of significance instead of attempting to find the true onset of changes in responses that must have occurred a little before. This criterion, along with the fact that the crossing of the threshold of 0.05 was followed by highly significant differences, ensured reliable conclusions. It should be noted that corrections for multiple comparisons do not apply here for two reasons: first the samples at each time step are involved in only one comparison, and second consecutive samples are not statistically independent. Indeed, if there is a significant difference at one time step, it is very likely that there is also a significant difference in the next time step because signals do not vary instantaneously. Hence, the risk of false positive must not be controlled.

An index of motor adaptation was derived based on the relationship between the lateral commanded force and the measured force along the same axis. Similar metrics were used previously (Crevecoeur et al., 2018), and were based on the fact that these correlations were sensitive to learning. Indeed, a perfect compensation for the force field would produce high correlation, and we documented previously that errors made by ignoring the robot dynamics were on the order of ~10%. Hence, a change in correlation from 0.4 to 0.7 or more on average with similar forward kinematics can be linked to adaptation. The data of the control experiment was also used to validate this argument empirically. Specifically, for each trial we computed a least-square linear regression between the commanded force obtained from velocity signals, and the applied force measured with the force encoders. These correlations were then averaged across perturbation directions for each participant (as they revealed qualitatively similar effects), then across participants for illustration. Surrogate correlations were obtained by calculating linear regressions between the measured force and the commanded force of randomly selected trials with replacement. These surrogate correlations were calculated on 100 randomly picked trials with replacement for each index and participant.

## Results

### Experiment 1

Our first experiment was designed to reproduce previous findings about adaptation of feedback responses to unpredictable disturbances, and measure accurately from EMG data the moment when the perturbation-related activity started to be tuned to the force field. Importantly, a feedback response is expected in all cases (Milner and Franklin, 2005; Wagner and Smith, 2008; Cluff and Scott, 2013). What we searched for was not just a feedback response, but a change in feedback response across early and late force field trials indicating that the response became adapted to the force field.

We measured a clear deviation in the lateral cursor displacement in the direction of the force field as expected since the perturbations could not be anticipated (Fig. 1b). Although the maximum hand displacement exhibited a small reduction across the first few trials, the exponential fits of this variable as a function of trial index did not display any significant curvature (*P* > 0.05 for both directions). In contrast, the maximum target overshoot exhibited a clear and highly significant exponential decay across trials (Fig. 1c, *P*<0.001). Hence participants initiated force field trials with a controller that would otherwise produce a straight reach path (black traces in Fig. 1b), resulting in a clear perturbation-related movement error, but then managed to improve their online correction across perturbation trials.

It was previously suggested that the reduction in target overshoot did not result from an increase in control gains or in the mechanical impedance of the limb. Instead, it resulted from a reduction in interaction forces at the handle (Crevecoeur et al., 2018). As a consequence, we expected to measure a reduction in muscle response to the perturbation. To observe this, we averaged EMG data across the first four and last four trials for each perturbation direction. Our rationale was that fewer than four trials would likely be too small a sample, whereas trials with indices 5 and more were already displayed larger adaptation in particular for CW perturbations, thereby reducing the size of the effect under investigation. As expected, we found a significant reduction in EMG responses to the perturbations (Fig. 2a-b). When traces were aligned to the position threshold and averaged across directions (see Methods), we found that the onset of drop in the time series of *P*-values occurred on average at 122ms following threshold (center of the 30ms bin, Fig. 2c-d). It could be observed that there was no clear difference across the first and last trials, which could have indicated the presence of systematic co-activation on average.

**Figure 2.**
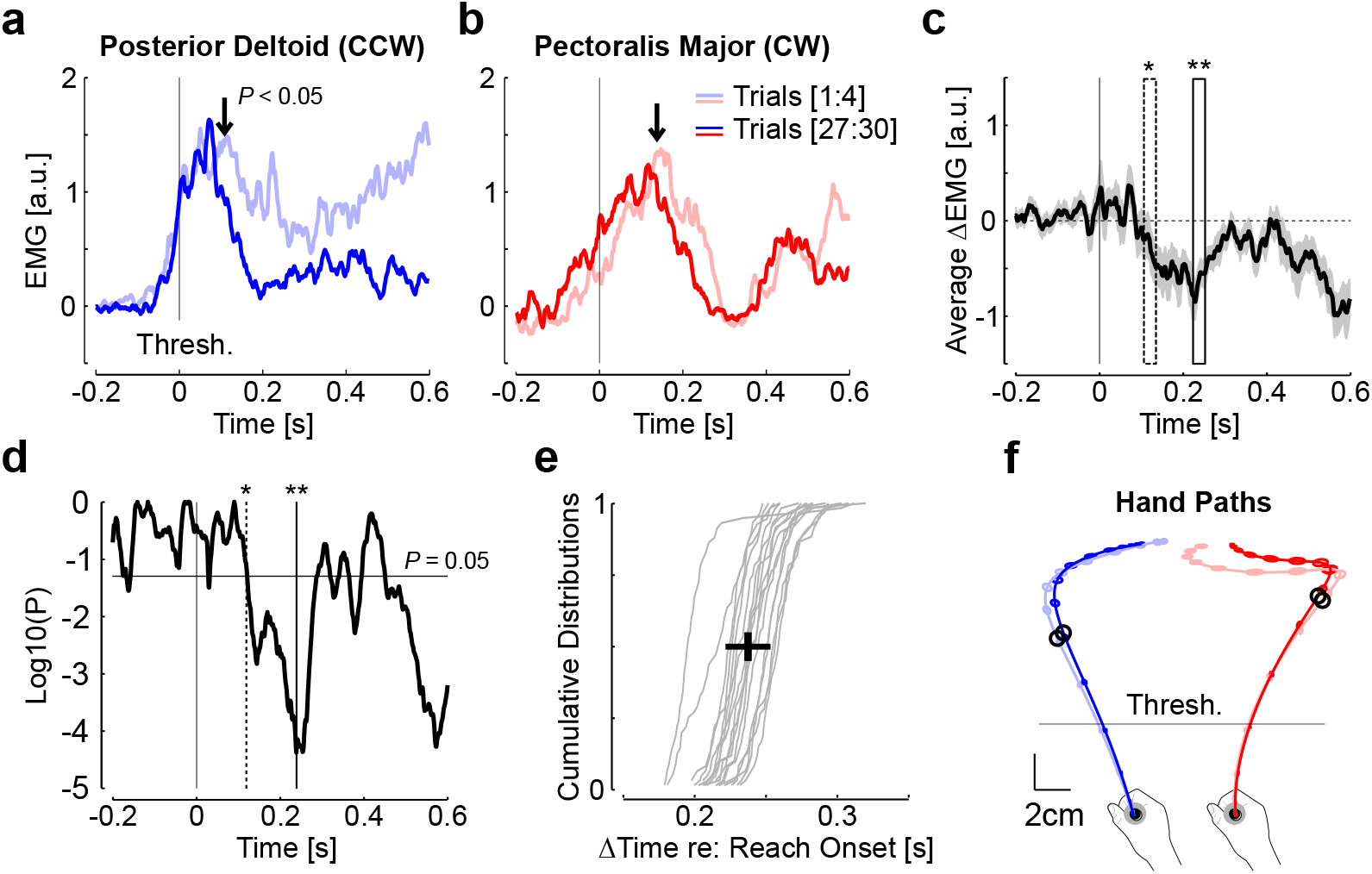
**a.** Activity of Posterior Deltoid averaged across the first four (light blue) and last four counter-clockwise (dark blue) perturbation trials. The vertical arrows illustrate the moment when a sliding paired comparison of the activity averaged in a 30m window dropped below P<0.05. Traces were aligned to the position threshold corresponding to one third of the reach path to reduce variability. **b.** Same as panel **a** for Pectoralis Major recorded from clockwise perturbation trials. The position threshold is also represented. **c.** Grand average of the difference between the activities of the first four and last four force field trials, aligned to the position threshold and averaged across muscles and participants (n=18). The gray area corresponds to the standard error of the mean. The dashed window is the first window that display significant difference from sliding paired comparison (P<0.05, width=30ms). The solid window is the window associated with the minimum P-value (P<10^−4^). **d.** P-value of the sliding paired comparison performed on the data from panel **c**. All EMG traces were smoothed with a 5-ms sliding window for illustration purposes. **e.** Cumulative distribution of the delay between movement onset, and the moment when the P-value of panel **d** dropped below 0.05. This moment corresponds to the time of threshold crossing plus 122ms. The median delay between movement onset and this time was 237±15ms (mean±SD across participants, n=18). **f.** Average hand paths and standard dispersion ellipses for the first four and last four trials in each direction, Dispersion ellipses are displayed 50ms (see Methods). The black dots represent the moment corresponding to the vertical arrows of panels **a** and **b**.

Because there was some variability between reach onset (defined as the moment when the cursor exited the home target) and the moment when participants’ hand crossed the position threshold, we calculated for each subject a distribution of elapsed time between reach onset and the moment corresponding to threshold + 122ms. These distributions are reported in Fig. 2e, and the mean±SD of medians is shown (black cross). The mean value was 237ms. For illustration, we reported in Fig. 2f the mean latency of within trial changes in feedback response on the average hand path represented for the first and last four trials. The black circle illustrates the moment of significant reduction in perturbation-related EMG that could be linked to the reduction in target overshoot observed in Figure 1.

The reduction in perturbation related response in EMG and in target overshoot were expected if participants learned to handle the force field. To further address whether their online corrections reflected adaptation, we correlated the measured lateral force with the commanded force calculated offline based on forward hand velocities. Average traces were represented in Fig. 3a for CCW perturbations (normalized for illustration). Observe that the average correction in the first trial was variable and the traces were irregular (Fig. 3c). In contrast, the same data plotted for the last trials appeared more regular (Fig. 3b, d). Figure 3c and 3d show a phase diagram with measured and commanded forces in the first and last trials for each participant. These traces were taken from ~200ms following reach onset to 1000ms (gray rectangles in panels a and b), based on the previous analysis revealing that there was no difference until ~240ms following reach onset, and thus no expected improvement in correlation prior to this time.

**Figure 3.**
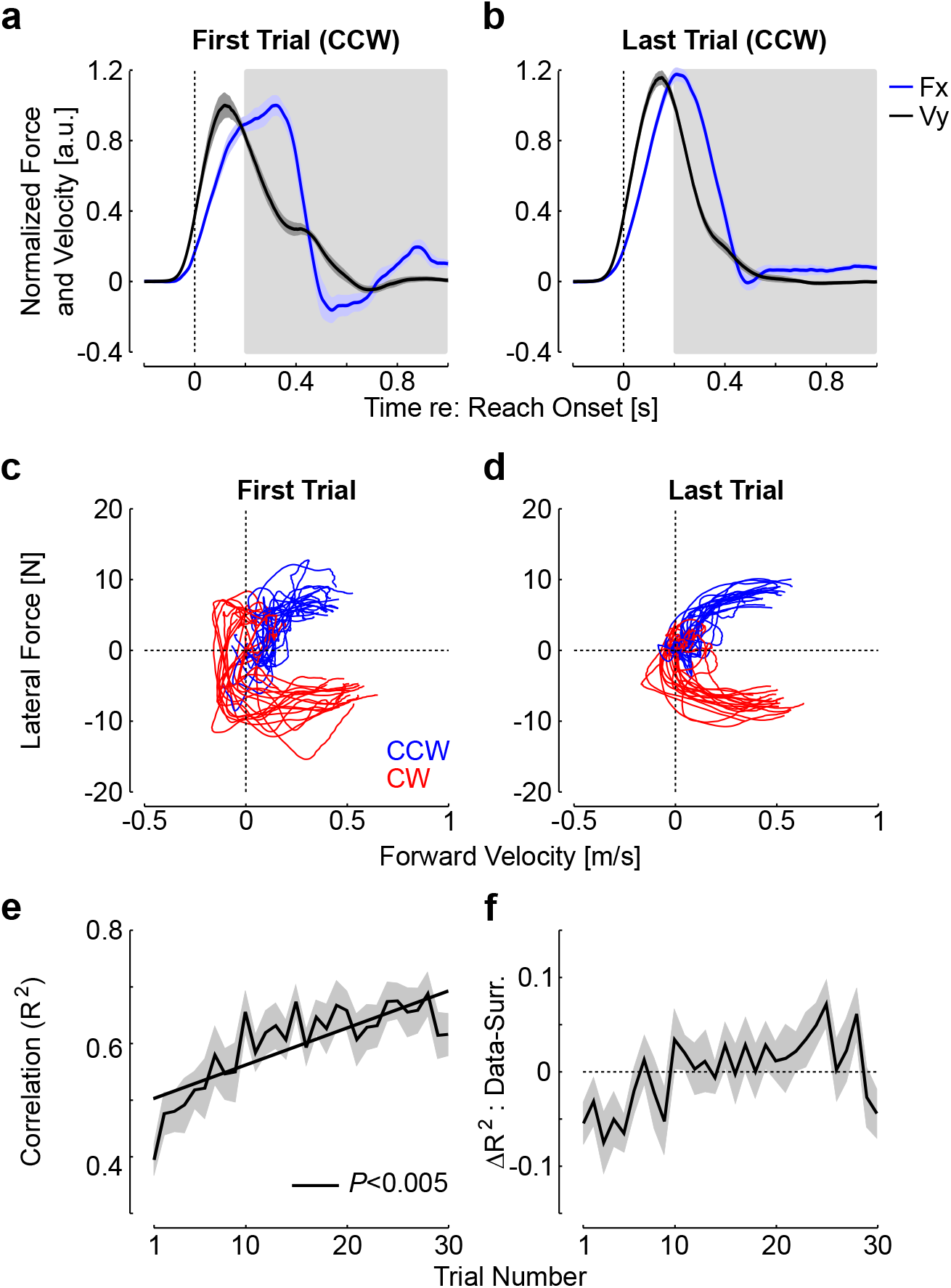
**a.** Forward hand velocity (black) and measured lateral force (blue) normalized to the average maximum calculated on the first trial. Shaded areas represent one standard error across participants (n=18). Panels display data from the first (left) and last (right) trials with counter-clockwise force field perturbation. **b.** Lateral force as a function of forward velocity from the first (left) and last (right) force field trials. Counter-clockwise and clockwise perturbations are shown in blue and red, respectively. Traces are from individual trials (one trace per participant). **c.** Solid: mean±SEM of the linear correlation (R^2^ statistics) across force field trials from Experiment 1 (CCW and CW perturbations averaged). Dashed: same for the first 30 trials from the control experiments for which perturbations were fully predictable (n=8, distinct group of participants). The orange lines illustrate the asymptote correlation ± SEM obtained by average the correlation across the last 50 trials from the control group. **d.** Difference between the correlations from Experiment 1 as calculated in panel c, and the correlation between the lateral force of each trials with the forward hand velocity of a randomly picked surrogate trial with replacement. The surrogate correlations were calculated 100 times per trial and participants, and averaged across. Shaded area is one SEM.

The correlations exhibited highly significant changes across trials. This was first assessed with a repeated measures ANOVA on the correlations with the trial indices as main factor (rmANOVA, *F*_(29,493)_= 8.7, *P*<10^−5^), and a standard least square linear regression highlighted a clear increase in this variable (Fig. 3e, *P*<0.005). These correlations were compared to those obtained with randomly picked surrogate profiles to see whether the measured force in each perturbation trial reflected tuning to the ongoing perturbations (see also Methods, Fig. 3f), or whether a non-specific correction pattern was produced, which could correlate as well with randomly picked surrogates as with the experienced perturbation. We found that the true correlations were initially below the surrogate, and then became greater than the surrogates. In support to this observation, we found a significant interaction between the correlation types (true versus surrogate) and the trial index (rmANOVA, *F*_(29,493)_=1.88, *P*=0.004). This analysis suggests that the force measured at the handle depended on the specific perturbation profile experienced during each force field trials.

We verified with the data from the control experiment that an increase in the same correlations between the measured and commanded forces occurred when the perturbations were fully predictable (Fig. 4). Likewise, the commanded and measured forces displayed initially variable traces with a terminal increase in interaction force at the handle consistent with the production of a target overshoot (black arrow), followed by a more regular and similar profiles. Hence, the key observations were that these correlations represented a sensitive metric of learning, and they increased across trials in the random context of Experiment 1 similarly as in the standard context of trial-by-trial adaptation. In all, the data from Experiment 1 highlighted that participants were able to adapt their feedback responses to unanticipated force field disturbances within ~240ms following reach onset. In addition, we found increases in correlations between applied and commanded forces that paralleled the behavior observed in a standard learning paradigm.

**Figure 4.**
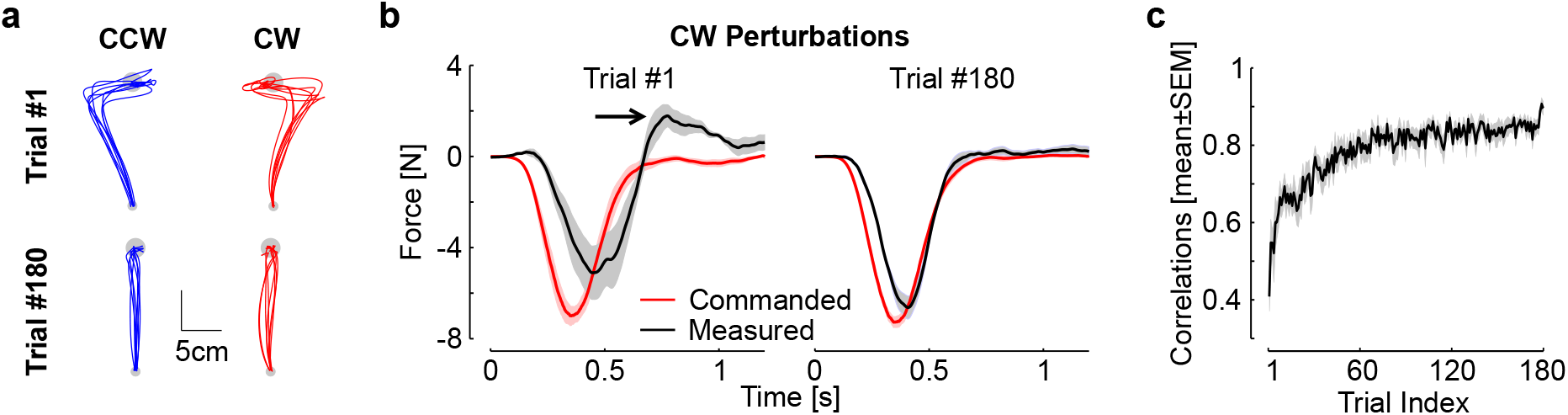
Control experiment, data from (Crevecoeur et al., 2018). **a.** Individual hand traces from the first and last perturbation trials in each direction from the control experiment (one traces per participant, n=8). **b.** Commanded (black) and measured (red) force profiles for the first (left) and last (right) clockwise perturbations. The arrow highlights the increase in peak terminal force linked with the target overshoot. Observe that the traces become very similar, which results in an increase in the temporal correlation between them. **c.** Correlations between commanded force and measured force as in Figure 3 against trial indices. Correlations were averaged across directions and participants. Displays are mean±SEM across participants.

### Experiment 2

This experiment was designed to investigate whether participants could learn to adapt their feedback responses when exposed to four different force fields at the same time: either orthogonal or curl force field, in clockwise or counter-clockwise directions. For the orthogonal field, we observed the same behavior as in Experiment 1: minute changes in the maximum lateral displacement across the first few trials, and highly significant exponential decay of the maximum target overshoot across trials (not shown). Hand traces during curl field trials were distinct because, unlike the orthogonal field, the antero-posterior component of the force field prevented a systematic target overshoot (Fig. 5a-b). For this reason, we used the path length to capture adaptation across the two force fields with the same metric. We found a clear reduction in path length for each force field and each perturbation direction (Fig. 5c). Exponential fits confirmed a very strong decay across trials (Fig. 5c, *P*<0.001).

**Figure 5.**
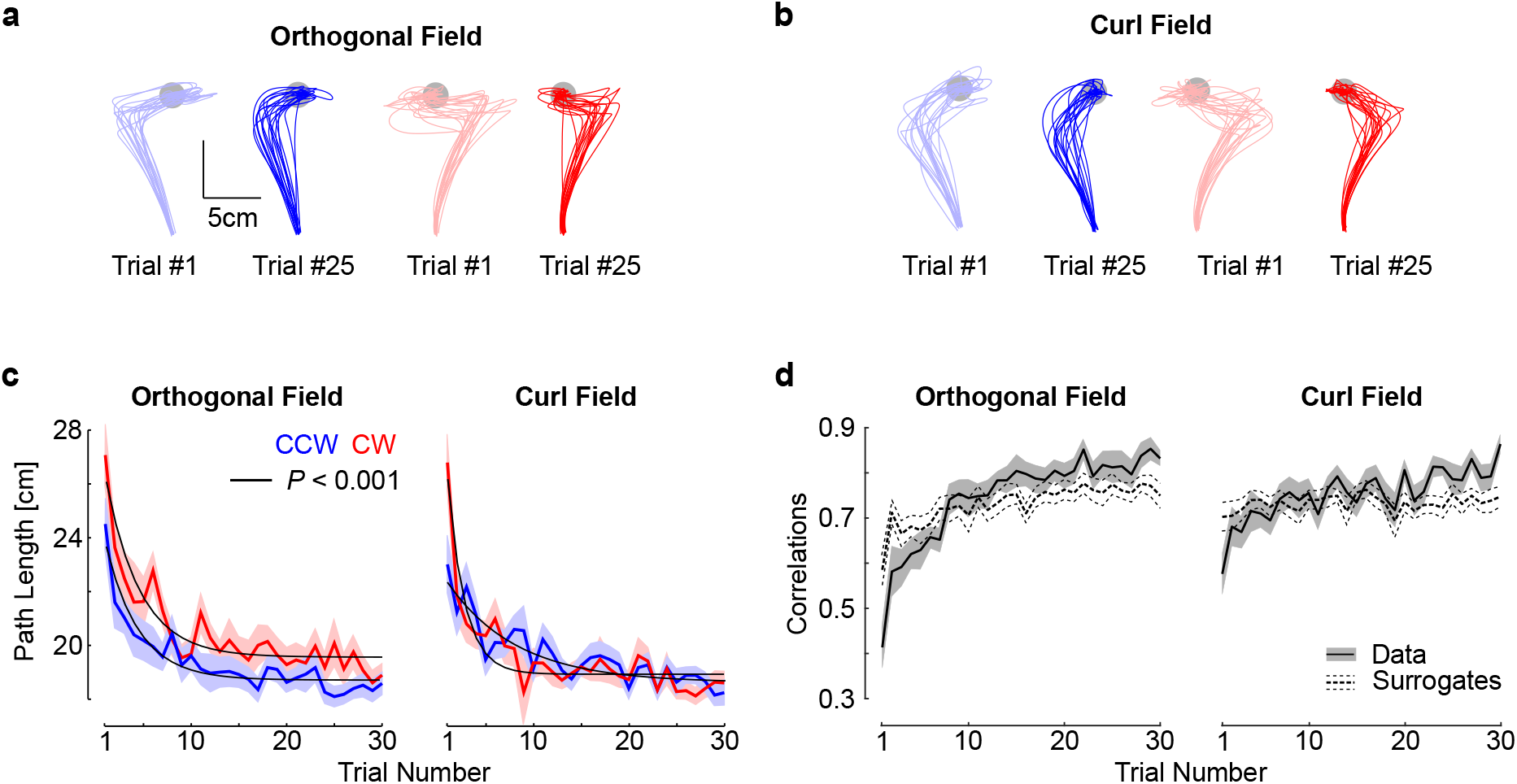
**a.** Hand paths from the first (light colors) and 25^th^ (dark colors) trials with an orthogonal field chosen to illustrate changes in feedback responses. All data were taken from Experiment 2, and each trace represent trials taken from each participant (n=18). Blue and red traces represent counter clock-wise and clockwise perturbations. **b.** Same as panel **a** for the curl field. **c.** Path length across force field trials. **d.** Trial by trial correlations between the lateral commanded force (proportional to forward velocity) and the measured force. Correlations were averaged across counter clockwise and clockwise directions. The shaded areas represent one SEM across participants. The dashed traces (mean±SEM) are the correlations between the measured lateral force and the commanded force corresponding to the velocity of randomly picked trials with replacement. The procedure was repeated 100 times for each participant, and the results were averaged across CCW and CW perturbations directions.

As for Experiment 1, we calculated the temporal correlations between the commanded and measured lateral forces averaged across clockwise and counterclockwise directions. For the two kinds of force fields, we found a clear impact of the trial index on the correlations, which confirmed the visible increase shown in Fig 5d (rmANOVA, *F*_(29,493)_>4, *P*<10^−6^). Furthermore, for the two kinds of force fields, we found highly significant interactions between the trial index, and the difference between the true and surrogate correlations (*F*_(29,493)_>5, *P*<10^−10^). This analysis indicated again that true correlations were lower first, then became greater, which supported that online corrections were tuned to the specific force profile experienced during each trial.

Surface recordings during the orthogonal force fields gave similar results as those reported in Fig. 2: we found that the initial responses to the perturbations were similar for Pectoralis Major and Posterior Deltoid, until 100ms following the threshold, where we observed a reduction in activity consistent with the production of an adapted response (Fig. 6a-c). As a consequence, the adjustments in this case were even observed slightly earlier than during Experiment 1. Indeed, the median time elapsed between reach onset, which was defined as the time when they exited the home target, and the reduction in target overshoot was 218±10ms (Fig. 6d). The analysis performed on curl field trials also revealed a significant reduction in perturbation-related activity occurring 104ms following the threshold, which corresponded to a delay between reach onset and changes in muscle response of 225±11ms (Fig. 6e-h).

**Figure 6.**
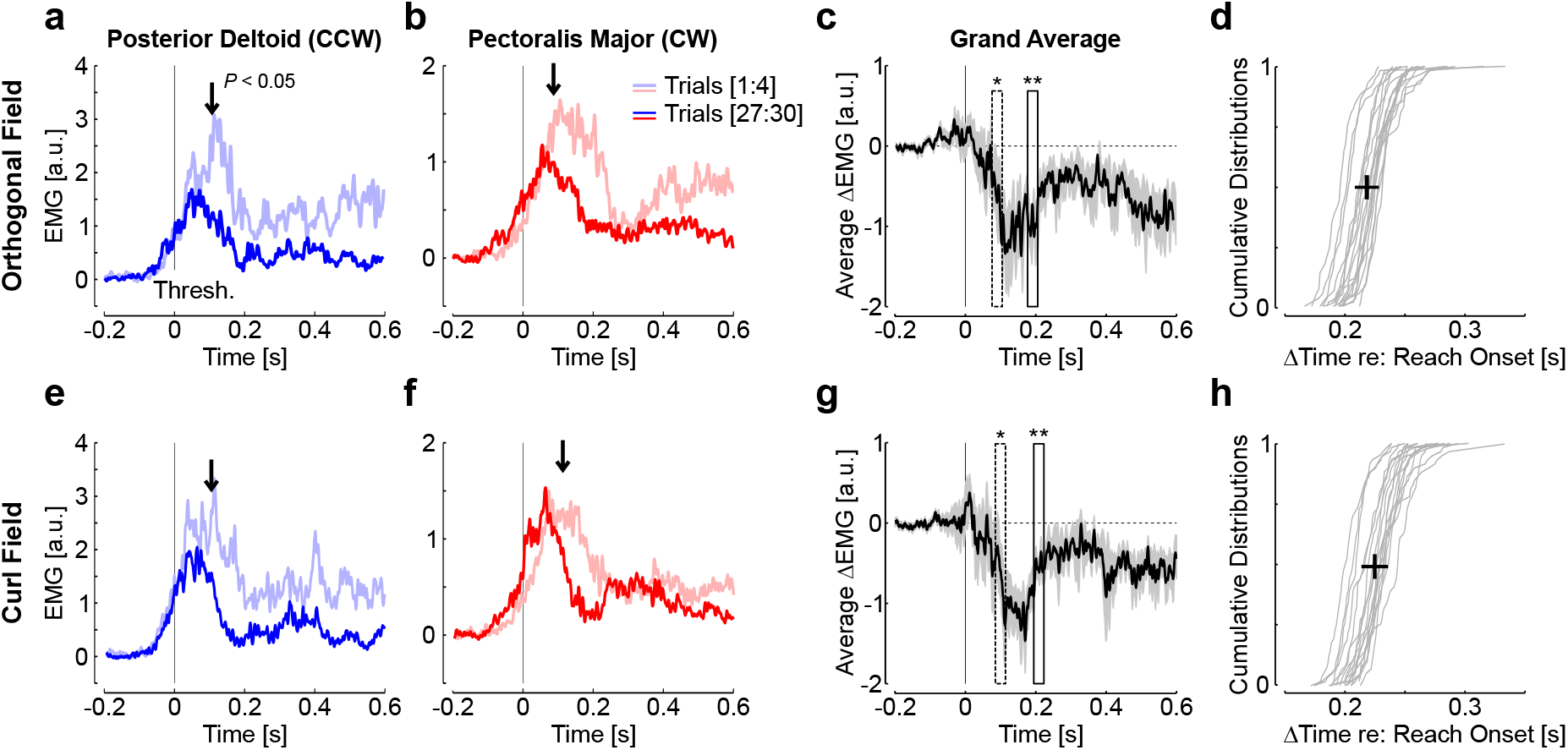
**a-d.** Same as Figure 2 for the orthogonal force field data from Experiment 2 (distinct group of 18 participants). Traces are first and last 4 trials in the force field for Pectoralis Major (**a**, blue) and Posterior Deltoid (**b**, red) aligned to the position threshold. The activity during unperturbed trials was subtracted and the traces were smoothed with a 5ms moving average for illustration. The vertical arrows show the moment with a sliding paired comparison of activity averaged in 30ms bins became significant (P<0.05). **c.** Difference between early and late feedback responses (mean±SEM) averaged across participants and muscles. The results of the sliding paired comparisons are shown: one star, first 30ms-bin with P<0.05, two stars: minimum of P (P<0.005, see Methods). **d**. Individual distribution of time elapsed between reach onset and the center of the first bin with P<0.05. The cross is the median ± SD across participants. **e-h.** Same as **a-d** for the curl force field.

Again, there was no systematic change in co-contraction, which could have impacted the mechanical impedance of the shoulder joint (Hogan, 1984; Burdet et al., 2001). Indeed, the average traces in Fig 6 c and 6g represent the average difference in muscle activity aligned to the position threshold each muscle. The presence of co-contraction would have resulted in an offset observed at the beginning of reach onset. To quantify this, we averaged the activity in each muscle from the first 100ms of the window presented in Fig. 6 (threshold-200ms until threshold-100ms), and compared the activity across the first and last four trials. We did not observe any statistical difference (t_(17)_<1.1, *P*>0.3). Besides possible changes in limb impedance (but see (Crevecoeur and Scott, 2014)), there is a known modulation of baseline activity evoked by unanticipated force field trials identified previously on the same muscles and during a similar task (Franklin et al., 2008; Crevecoeur et al., 2019). However, this effect did not impact the activity prior to the perturbations systematically, likely due to the fact that perturbation trials were randomly interspersed and the modulation of baseline activity averaged out.

It remained to be elucidated whether the feedback responses to the orthogonal and curl force fields were distinct, or whether participants used a single response pattern undifferentiated across force field disturbance, should the perturbations be sufficiently close to be handled with a single and non-specific response. Our data allowed us to reject this possibility. Indeed, we contrasted the feedback response to curl and orthogonal force fields averaged across the last 15 force field trials for each muscle. We found clear changes in EMG patterns, such that the curl force field evoked a stronger response, and was followed by a second increase in activity near the end of the reach (Fig. 7a-b). Based on the difference between the activities from curl and orthogonal fields, we found that the modulation of EMG activity was highly significant (Fig. 7c). Furthermore, the onset of changes in EMG revealed that the feedback responses were tuned to the force field very early during movement: this time corresponded to ~55ms prior to threshold on average (Fig. 7d), which was denoted as a star in the average hand path displayed in Fig. 7d (black circle). This median of the distributions of elapsed time from reach onset to this time across participants was 65±10ms (mean±SD).

**Figure 7.**
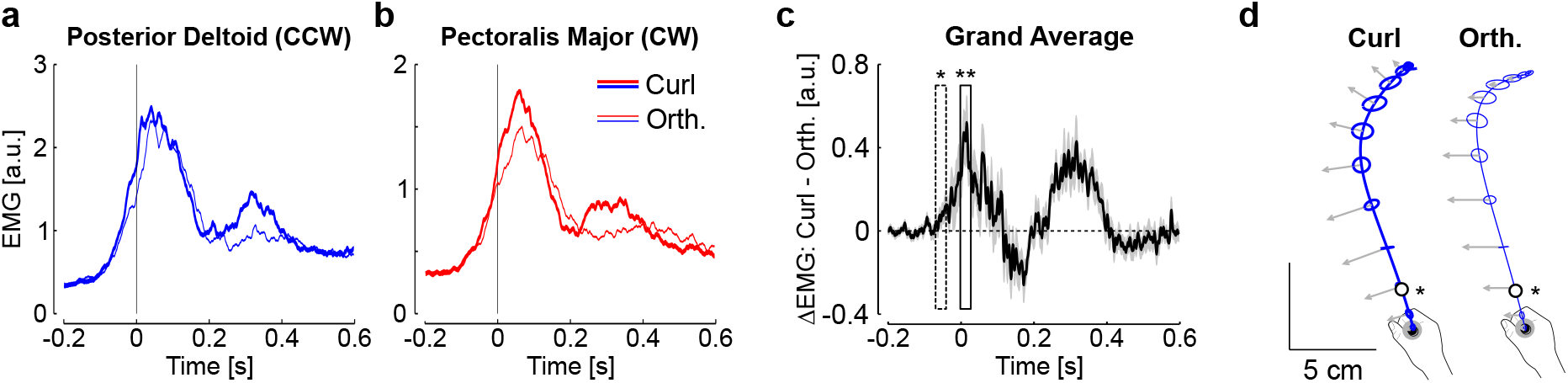
**a.** Average activity of Posterior Deltoid in curl (thick) and orthogonal (thin) counter clockwise perturbations across the last 15 trials of each type of force field. **b.** Same as **a** for Pectoralis Major from clockwise perturbations. **c**. Difference between activities recorded during curl and orthogonal force fields averaged across participants and muscles. The onset of significant changes based on a 30ms wide sliding window is highlighted with one star (P<0.05), followed by strongly significant differences (two stars, P<0.005). **d.** Average hand paths during counter-clockwise perturbations. Ellipses are 2-dimensional standard dispersion across participants every 50ms, and the impact of the force field is illustrated with gray arrow (a common scaling for the two force fields was applied for illustration). The open dots are the moment when the activities started to differ across to two types of perturbations.

To summarize, participants produced feedback responses tailored to the details of the unanticipated perturbations for each kind of force field; these feedback responses improved and exhibited similar traits as those of standard adaptation paradigms, namely the increase in correlation between commanded and measured forces. These changes in behavior were linked to finely tuned EMG activities, which indicated that feedback response was adapted to the force field as early as 250ms following reach onset.

## Discussion

Current theories of motor learning have postulated that sensory feedback about movement error is mapped to model updates for the next movement. Based on this idea, several seminal studies have characterized learning across movements by means of trial-by-trial learning curves. We recently argued that motor adaptation unfolded over faster time scales, potentially within a single trial, which revealed a novel function of motor adaptation, that is to complement feedback control online (Crevecoeur et al., 2018). To further test this hypothesis, we performed here two experiments with the following aims: 1- to reproduce our previous results on improvements in feedback responses to unanticipated force fields, 2- to identify in muscle recordings the latency of adaptive changes in control, and 3- to test whether healthy humans could learn two different force fields and two different perturbation directions at the same time. The results confirmed our previous findings, and highlighted that feedback responses became adapted within 250ms of reach onset. Importantly, feedback responses were also specific to each perturbation profile (curl and orthogonal force fields.

Our reasoning was based on the theory of adaptive control. The basic premise of this theory is that the controller adjusts the parameters of the state-feedback control loop in real time. In principle, there is no lower bound on the time scale of this mechanism, but the instantaneous learning rate should not be too large to prevent instability (Bitmead et al., 1990). The reduction in target overshoot for the orthogonal field (data from Exp. 1) without impacting the maximum lateral hand deviation was previously explained in this context (Crevecoeur et al., 2018). The ability to learn two different force fields at the same time was also a prediction of this model: if parameters can be tracked online, it does not matter which force field is applied (curl or orthogonal), since sensory feedback of each specific trial can be used to produce a response adapted to the ongoing perturbation.

Our results contrast with previous work reporting limited or no adaptation to different force fields that were cued prior to reach execution (Hirashima and Nozaki, 2012; Sheahan et al., 2016). In these studies, participants received explicit cues about whether the perturbation would be clockwise or counterclockwise force field. It was concluded that participants were not able to learn in comparison with other conditions such as when distinct representations (Hirashima and Nozaki, 2012), or distinct planning of a follow through movement (Sheahan et al., 2016) were associated with reaching. We did not test whether the performance in our experiment could be improved with explicit cues about the force field. Yet, it is clear that similar to these previous studies, we would not confirm the presence of adaptation based on measurements such as the maximum lateral displacement alone. The main difference is that we documented small changes in EMG and behavior, limited by movement duration and neural processing delays, but these changes were reliable and clearly reflected adaptation of feedback control. We do not question the fact that learning in a random paradigm is limited, but our data highlighted that motor execution can form and recall distinct internal representations of dynamics during movement.

Another candidate mechanism to counter unexpected disturbances is the use of co-contraction to modulate the limb’s intrinsic properties (Hogan, 1984; Burdet et al., 2001; Franklin et al., 2008), and the feedback gains in a non-specific way (Crevecoeur et al., 2019). Such strategy was previously demonstrated to influence feedback control gains across a few trials, which also limits the extent of lateral displacement following disturbances. However, our current dataset requires another explanation because the perturbations were applied randomly, and there was no co-contraction observed on average. Furthermore, an increase in EMG activity expected if a strategy based on co-contraction were used, whereas we documented a decrease in perturbation-related response across the two experiments (as shown in Fig. 2a, b, 6a, b, e, f). The absence of baseline co-activation questioned the possibility that a default increase in intrinsic impedance or in control gains were responsible for the adjustments of control.

Online tracking of model parameters is a candidate model to explain the change in control occurring within a movement. Indeed, movements were straight on average and force field trials were often separated by several baseline trials. Thus, participants initiated the reaching movements with a controller corresponding to a baseline trial (i.e. without force field), which would in this case produce a straight reach path (see black traces in Fig. 1). Then, during perturbations, they changed their control to produce feedback responses that became adapted to the force field. This transition between a baseline controller and a controller adjusted to each force field, along with the observation that each feedback response was better adapted without practicing in a predictable context, constituted a strong evidence for adaptive control in the motor system.

How much the controller changed within perturbation trials, or between two trials remains a matter of debate. On the one hand, our previous study provided an upper bound of ~500ms within which after effects could be evoked (Crevecoeur et al., 2018). Our current measurements based on EMG indicated that the change in feedback responses, likely based on the same mechanism, occurred within 250ms. This time window leaves enough room for adjustments of the controller to each force field within movement. However, it is clear that changes in movement representation also occurred offline, between two trials or over longer timescales (Krakauer and Shadmehr, 2006; Smith et al.; Kording et al., 2007; Dayan and Cohen, 2011).

Further investigations are required to better characterize the components of adaptive control. For instance, our experiments did not allow teasing apart how much vision and somatosensory feedback contributed to feedback adaptation, as the cursor was visible all the time. However, a strong contribution of proprioception is expected: first this system could produce detailed responses to the smallest perturbations with long-latency delays (50-60ms) (Crevecoeur et al., 2012), which almost certainly contributed to the early changes in EMG data of Experiment 2, evoked by very small differences in the force components across force fields. Furthermore, previous work highlighted that long-latency feedback engaged motor responses that are well captured in a state-feedback control model (Crevecoeur and Kurtzer, 2018). This rapid state-feedback control loop is supported by a distributed network through primary sensory and motor cortices, pre-motor cortex, parietal regions, and cerebellum (Flament et al., 1984; Omrani et al., 2016). Hence, the fastest adjustments to state feedback control could be achieved by tuning the long-latency feedback loop.

Besides the potential contribution of muscle afferent feedback, the fact that changes in feedback responses were detected within 250ms leaves enough time to engage task-related feedback responses mediated by touch (Pruszynski et al., 2016; Crevecoeur et al., 2017), and vision, which participates in goal-directed feedback control (Franklin and Wolpert, 2008; Scott, 2016), as well as in rapid changes in navigation strategies (Cross et al., 2019). Characterizing the specific contribution of each sensory system constitutes an exciting challenge for future work.

